# The Trials and Tribulations of Priors and Posteriors in Bayesian Timing of Divergence Analyses: the Age of Butterflies Revisited

**DOI:** 10.1101/259184

**Authors:** Nicolas Chazot, Niklas Wahlberg, André Victor Lucci Freitas, Charles Mitter, Conrad Labandeira, Jae-Cheon Sohn, Ranjit Kumar Sahoo, Noemy Seraphim, Rienk De Jong, Maria Heikkilä

## Abstract

The need for robust estimates of times of divergence is essential for downstream analyses, yet assessing this robustness is still rare. We generated a time-calibrated genus-level phylogeny of butterflies (Papilionoidea), including 994 taxa, up to 10 gene fragments and an unprecedented set of 12 fossils and 10 host-plant node calibration points. We compared marginal priors and posterior distributions to assess the relative importance of the former on the latter. This approach revealed a strong influence of the set of priors on the root age but for most calibrated nodes posterior distributions shifted from the marginal prior, indicating significant information in the molecular dataset. We also tested the effects of changing assumptions for fossil calibration priors and the tree prior. Using a very conservative approach we estimated an origin of butterflies at 107.6 Ma, approximately equivalent to the Early Cretaceous–Late Cretaceous boundary, with a credibility interval ranging from 89.5 Ma (mid Late Cretaceous) to 129.5 Ma (mid Early Cretaceous). This estimate was robust to alternative analyses changing core assumptions. With 994 genera, this tree provides a comprehensive source of secondary calibrations for studies on butterflies.

## INTRODUCTION

An increasing amount of molecular information is allowing the inference of broad and densely sampled phylogenetic hypotheses for species-rich groups. This effort, combined with the emergence of a great number of methods investigating trait evolution, historical biogeography, and the dynamics of diversification have increased the need for time-calibrated trees. Estimating divergence times in molecular phylogenetic work depends primarily on fossils to constrain models of heterogeneous rates of substitutions. Consequently, the robustness of such estimates relies on the quality of fossil information, involving age and taxonomic assignment (Parham et al 2012), the priors assigned to nodes that are calibrated in a Bayesian analysis (Warnock et al 2012, Brown & Smith 2017), and the amount of information inherent in the molecular dataset (Yang & Rannala 2006, Rannala & Yang 2007, dos Reis & Yang 2013).

Fossils inform us of the minimum age of a divergence, imposing a temporal constraint that is widely accepted. However, the constraint of a simple hard minimum age is insufficient information for a proper analysis of times of divergence, particularly as there is an absence of information about maximum ages for divergences, including the root node. Often fossil information is modeled as a probability distribution, such as a lognormal or exponential distribution, indicating our beliefs regarding how informative a fossil is about the age of a divergence (Drummond et al 2006, Warnock et al. 2015). The distributional shapes of these priors are often established without justification (Warnock et al. 2012). Ideally, in node-based dating, fossil information is used only as a minimum age constraint for a given divergence in the form of a uniform prior with a minimum age equaling the fossil age and a maximum age extending beyond the age of the clade in question. In such cases at least one maximum constraint is needed, often also based on fossil information. Another approach is use of extraneous additional information, such as using ages of host plant families as maximum constraints for highly specialized phytophagous insect clades (Wahlberg et al. 2009). In such cases, a uniform prior also can be used, with the maximum set to the age of the divergence of the host plant family from its sister group and the minimum set to the present time.

Brown & Smith (2017) recently have pointed out the importance of assessing the relative influence of priors over the actual amount of information contained in the molecular dataset. As noted above, users specify fossil calibrations using prior distributions by modeling the prior expectation about the age of the node constrained. However, the broader set of fossil constraints can interact with each other and with the tree prior, leading to marginal prior distributions at nodes that usually differ from the user’s first intention (Warnock et al 2012). If relevant information were contained within the molecular dataset, one would expect the posterior distribution to shift from the marginal prior distribution. In the case of angiospermous plants, Brown & Smith (2017) showed that the marginal prior resulting from the interaction of all priors (fossils and the tree) excluded an Early Cretaceous origin, in effect giving such an origin zero probability. In addition, many calibrated internal nodes showed nearly complete overlap of marginal prior and posterior distributions, suggesting little information in the molecular dataset but a potentially strong influence of the set of priors.

With more than 18,000 species described and extraordinary efforts made to infer phylogenetic hypotheses based on molecular data, butterflies (Lepidoptera: Papilionoidea) have become a model system for insect diversification studies. Nevertheless, the paucity of information available to infer times of divergence in butterflies questions the reliability of the various estimates (e.g. Garzón-Orduña et al. 2015). Heikkilä et al. (2012) for example, used only three fossils to calibrate a higher-level phylogeny of the superfamily Papilionoidea. The shortage of fossil information for calibrating large-scale phylogenies also means that, most of the time, species-level phylogenies at a smaller scale rely on secondary calibration points extracted from the higher-level time-trees (e.g. Peña et al. 2011, Matos-Maravi et al. 2013, Kozak et al. 2015, Chazot et al. 2016, Toussaint & Balke 2016).

In a recent paper, de Jong (2017) revisited the butterfly fossil record, providing a discussion about the quality of the different fossil specimens as well as their taxonomic placement. Using this information, we established an unprecedented set of 12 fossil calibration points across all butterflies, which we use in this study to revisit the timescale of butterfly evolution in a comprehensive phylogenetic framework, and investigate the robustness of this new estimate. We complement the minimum age constraints of clades based on fossils with maximum age constraints based on the ages of host plant families. Some clades of butterflies have specialized on specific groups of angiosperm hosts for larval development, such that one may assume that diversification of the associated butterfly clade only occurred after the appearance of the host plant clade. We use this assumption as additional information to calibrate the molecular clock by setting the age of specific clades of butterflies to be younger than the estimated age of their host plant lineage. We restrained these calibrations to higher-level host plant clades.

The most recent estimates of divergence times using representatives of all butterfly families inferred a crown clade age of butterflies of 110 Ma (Heikkilä et al. 2012) and 104 Ma (Wahlberg et al. 2013). These two dates yield to a large discrepancy when taking the fossil record into account, as the oldest known fossil butterfly is estimated to be 55.6 Ma and can be confidently assigned to the extant family Hesperiidae (de Jong 2016, 2017). Such discrepancy has been extensively debated for a similar case, the origin of angiosperms, often estimated to have originated during the Triassic (252–201 Ma ago), while the oldest undisputed fossil is pollen dated at 136 Ma. Despite a much more fragmentary fossil record for butterflies, the same questions remain. First, are the previous estimates robust to a more comprehensive assemblage of fossils and taxon sampling? Second, is the 50 million-year discrepancy between molecular clock estimates and the fossil record accurate or the result of a lack of information contained in the molecular dataset? In other words, how much does the set of priors influence the results?

Here, we generated a genus-level phylogeny of Papilionoidea, including 994 taxa, in order to maximize the number and position of fossil calibration points and increase the potential amount of molecular information. By establishing the set of 12 fossils and 10 host-plant calibration points, we time-calibrated the tree in order to provide a revised estimate of the timing in diversification of butterflies. We then assessed the robustness of these results to the assumptions made throughout the analysis, including (i) different subsets of fossil constraints, (ii) the prior distributions of fossil constraints, (iii) a different estimate for host plant ages, (iv) a Yule tree prior, (v) a reduced taxon sampling and (vi) the addition of a mitochondrial gene fragment to the nine nuclear gene regions.

Finally, we compared the user specified priors, marginal prior and posterior distributions of different analyses, to assess the influence of our set of constraints on the estimated timing of divergence.

## MATERIALS AND METHODS

### Molecular Dataset

When designing our dataset, we aimed at building a genus-level tree of Papilionoidea. We assembled a dataset of 994 taxa from the database VoSeq (http://www.nymphalidae.net/db.php, Peña & Malm 2012), with each taxon representing a genus. We chose to include gene fragments that were available across the whole tree in order to avoid large clade-specific gaps in the molecular dataset. In addition, Sahoo et al. (2016) pointed out a conflicting signal in the family Hesperiidae between nuclear and mitochondrial markers. Thus, we chose to primarily focus on nuclear markers. Our final dataset included nine gene fragments: ArgKin (596bp), CAD (850bp), EFI-□ (1240 bp), GAPDH (691bp), IDH (710 bp), MDH (733 bp), RPS2 (411 bp), RPS5 (617 bp) and wingless (412 bp) for a total length of 6260 base pairs. The list of taxa and Genbank accession codes are available in the Supplementary Material S1.

### Set of Time-Calibrations for Timing Analyses

Fossil calibrations – Previous studies estimating times of divergence of butterfly lineages have largely relied on unverified fossil calibrations. The identifications of these calibrations were often based on overall similarity with extant taxa, not apomorphies. In the present study, we initially chose 14 fossil butterflies that were recently critically reviewed by de Jong (2017) and displayed apomorphic characters or character combinations diagnostic of extant clades, thereby allowing reliable allocation of fossils on the phylogenetic tree to provide minimum ages to the corresponding nodes. These fossils included three inclusions in Dominican Amber and 11 compression/impression fossils. For the age of these fossils we have relied on the most recent dates established from recent advances in Cenozoic chronostragigraphy, geochronology, chemostratigraphy and the geomagnetic polarity time scale (Walker et al., 2013). These improvements by geologists and specialists in allied disciplines have provided an increased precision in age dates of stratigraphic record (International Commission on Stratigraphy, 2012). The list of fossils and their positions in the tree is given in Table 1 and Figure 1. For more detailed information on the identification of these fossils, localities, preservation type and current depositories, see de Jong (2017).

**Figure 1.**
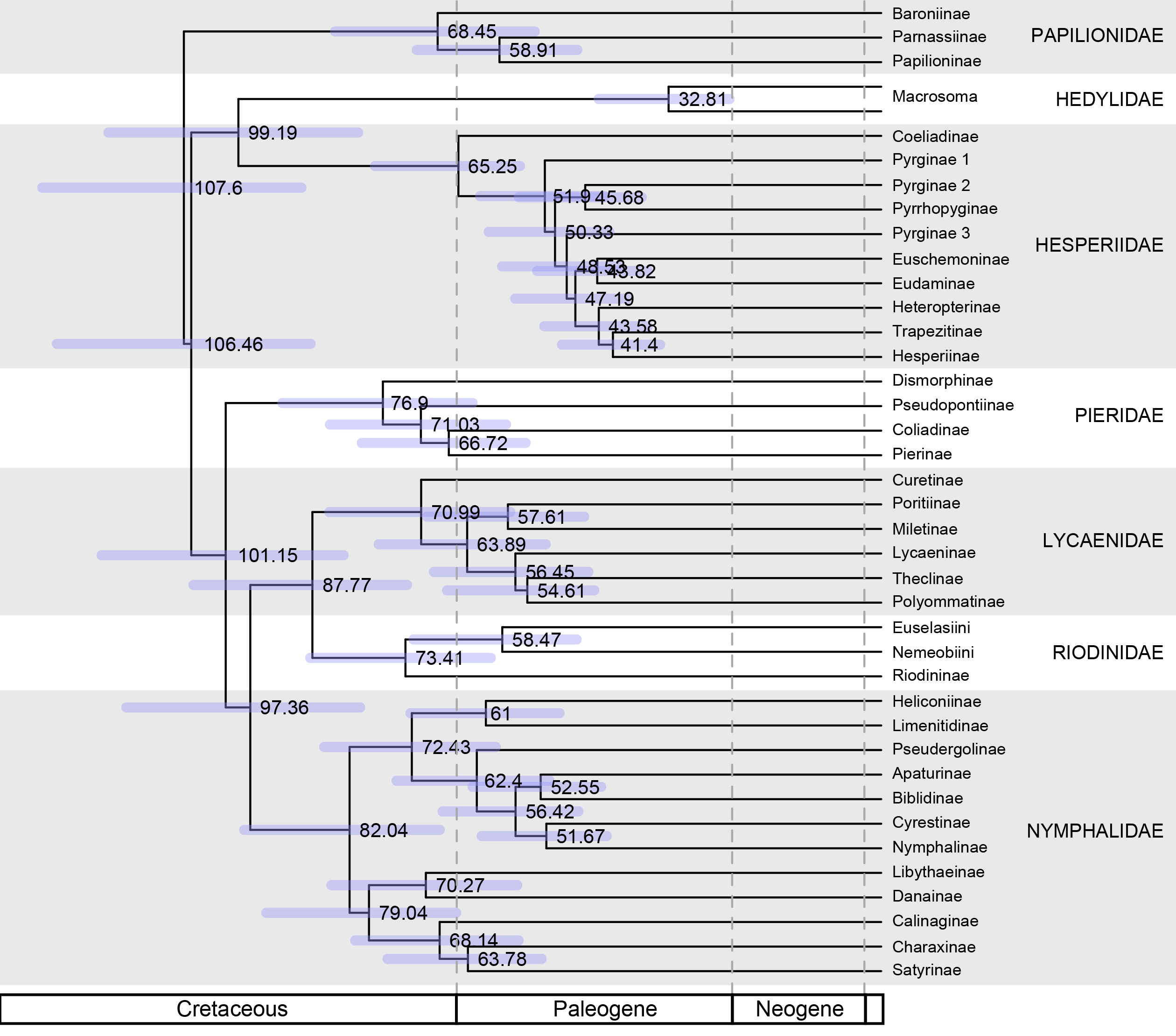
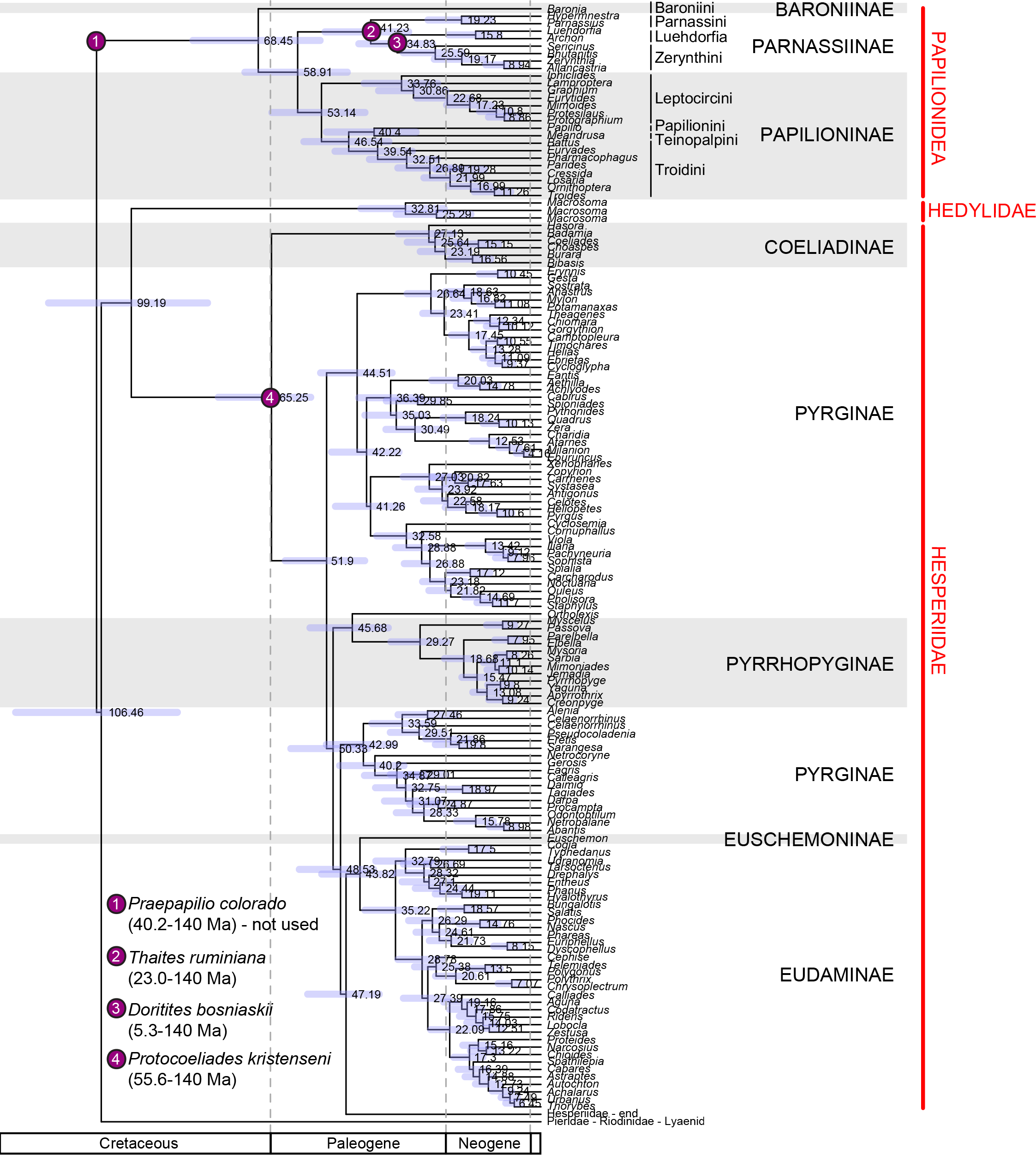
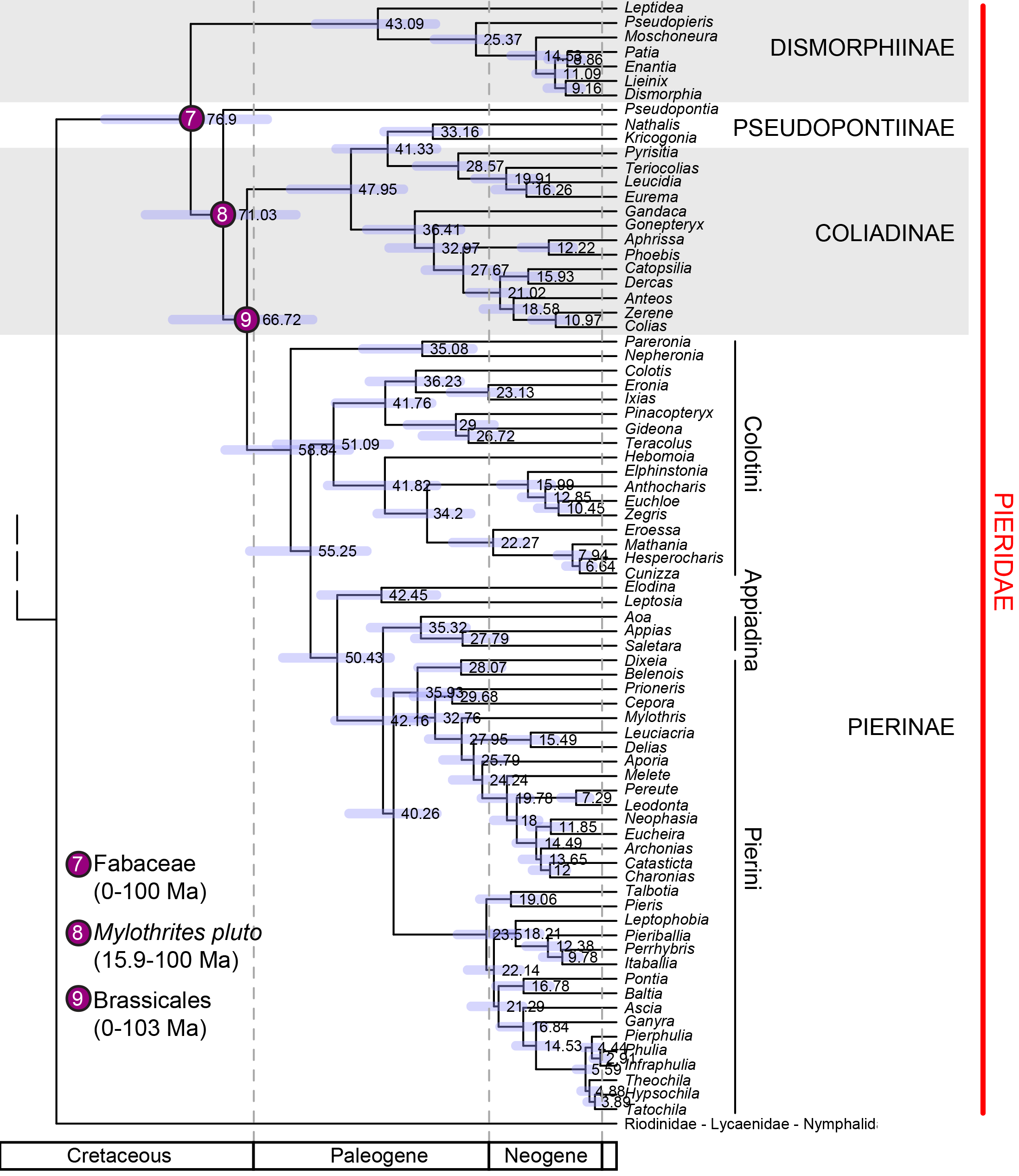
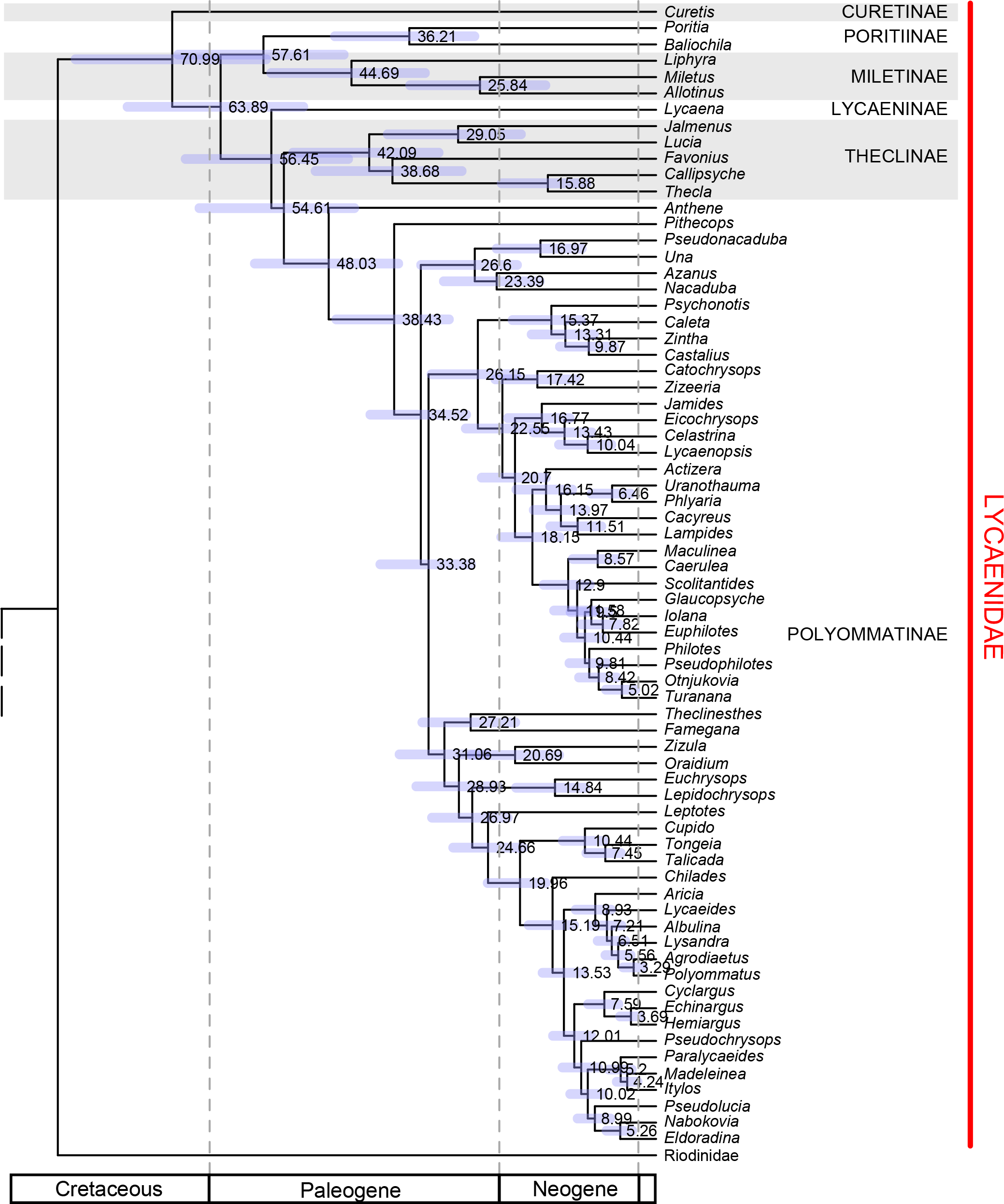
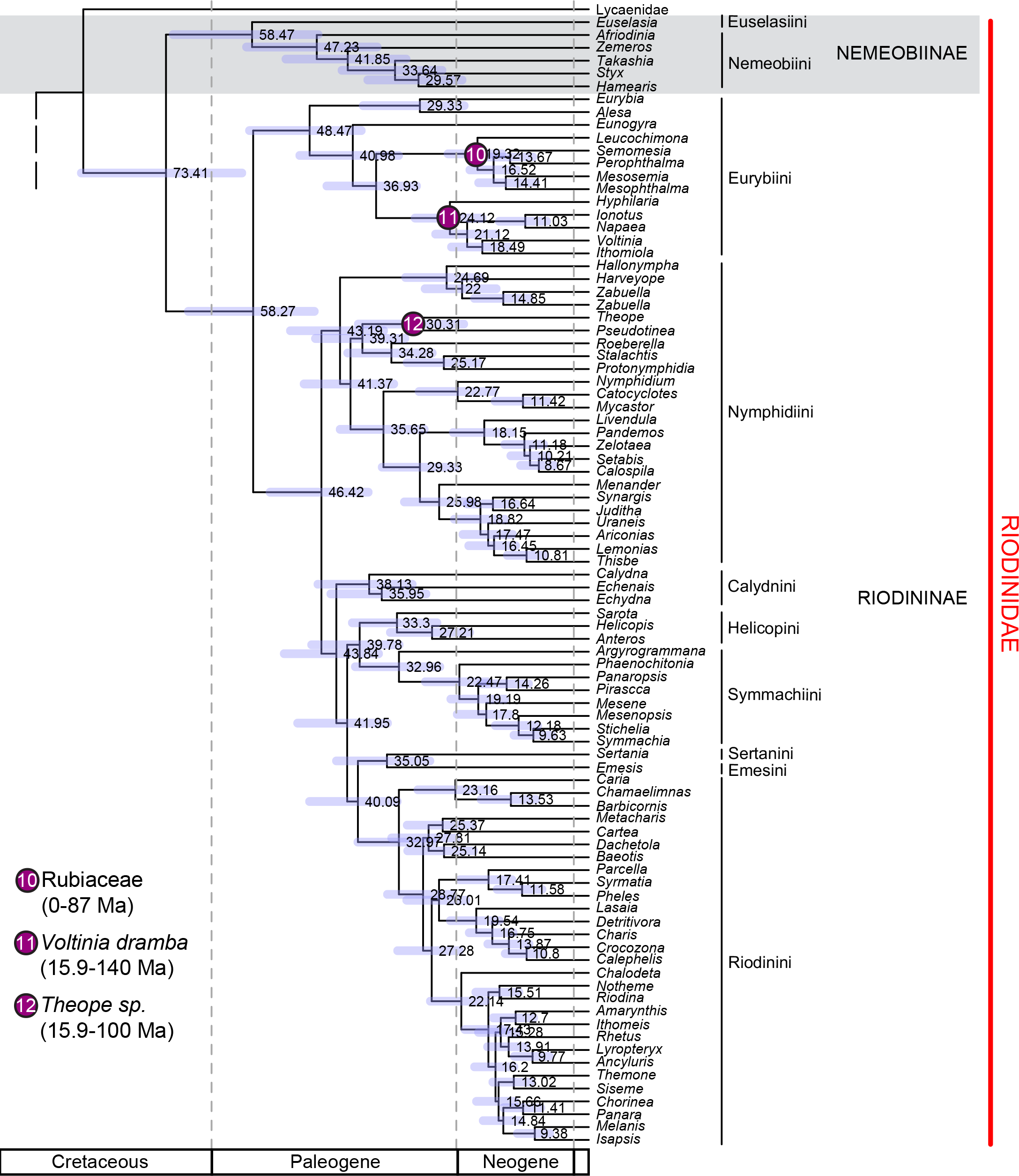
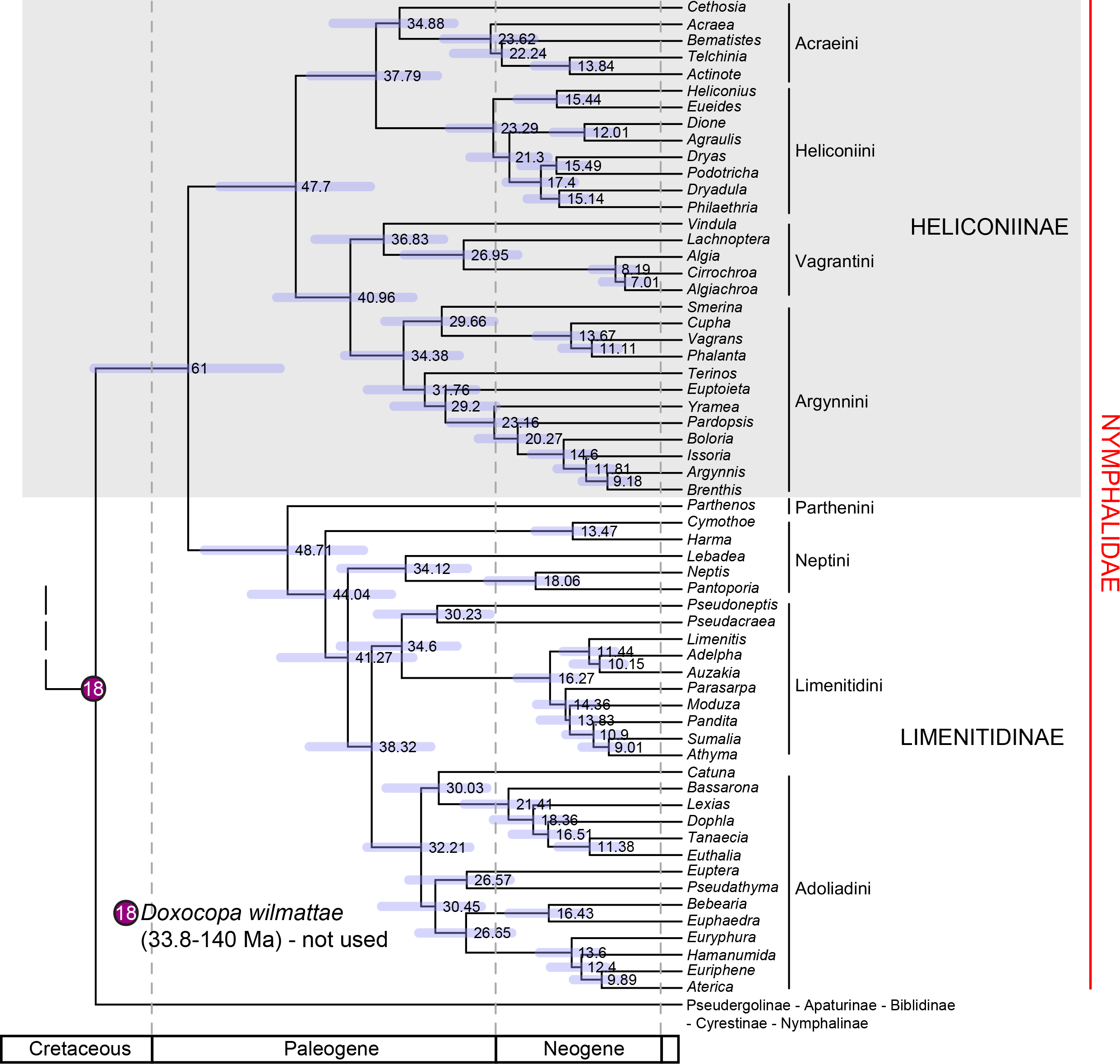
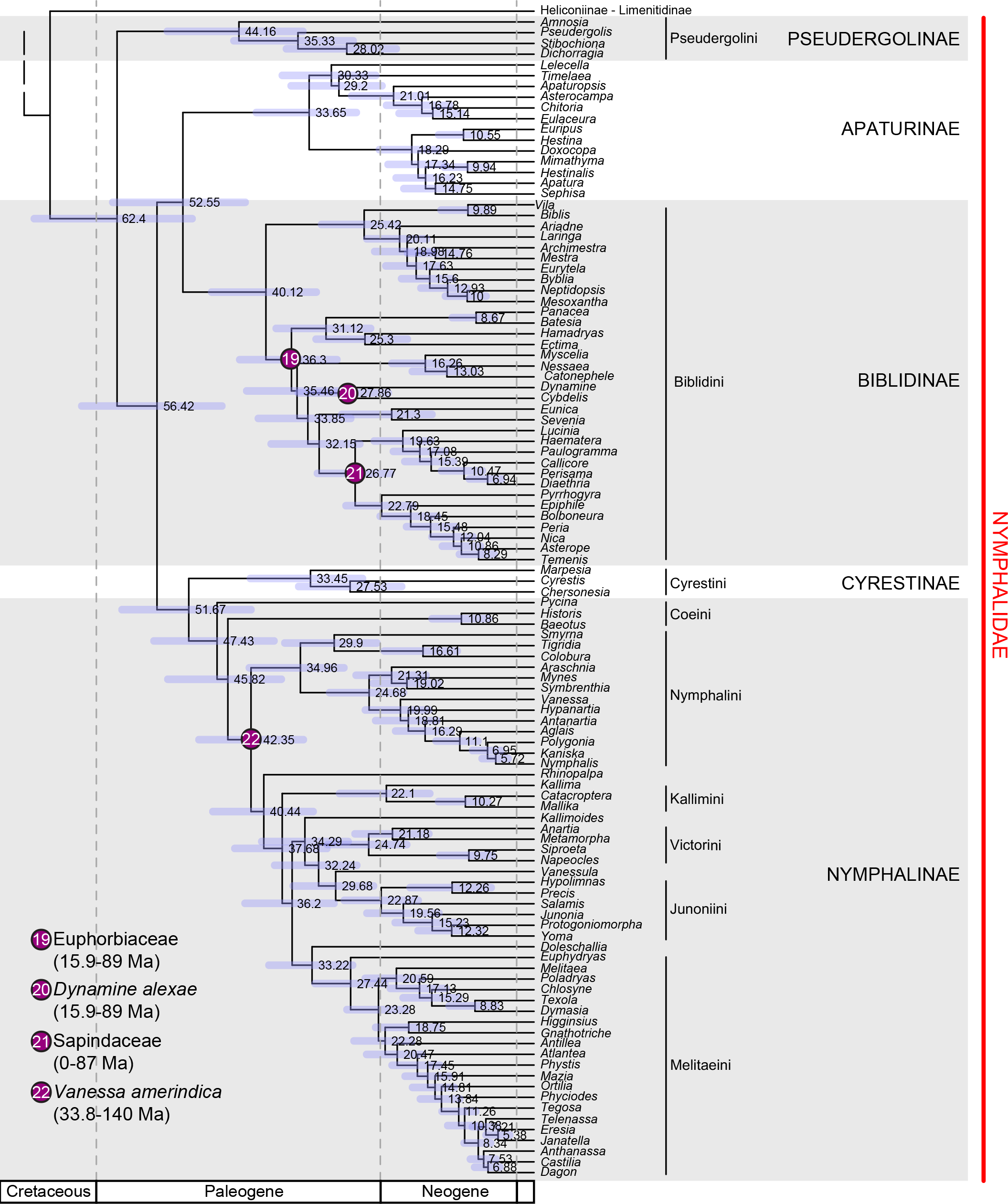
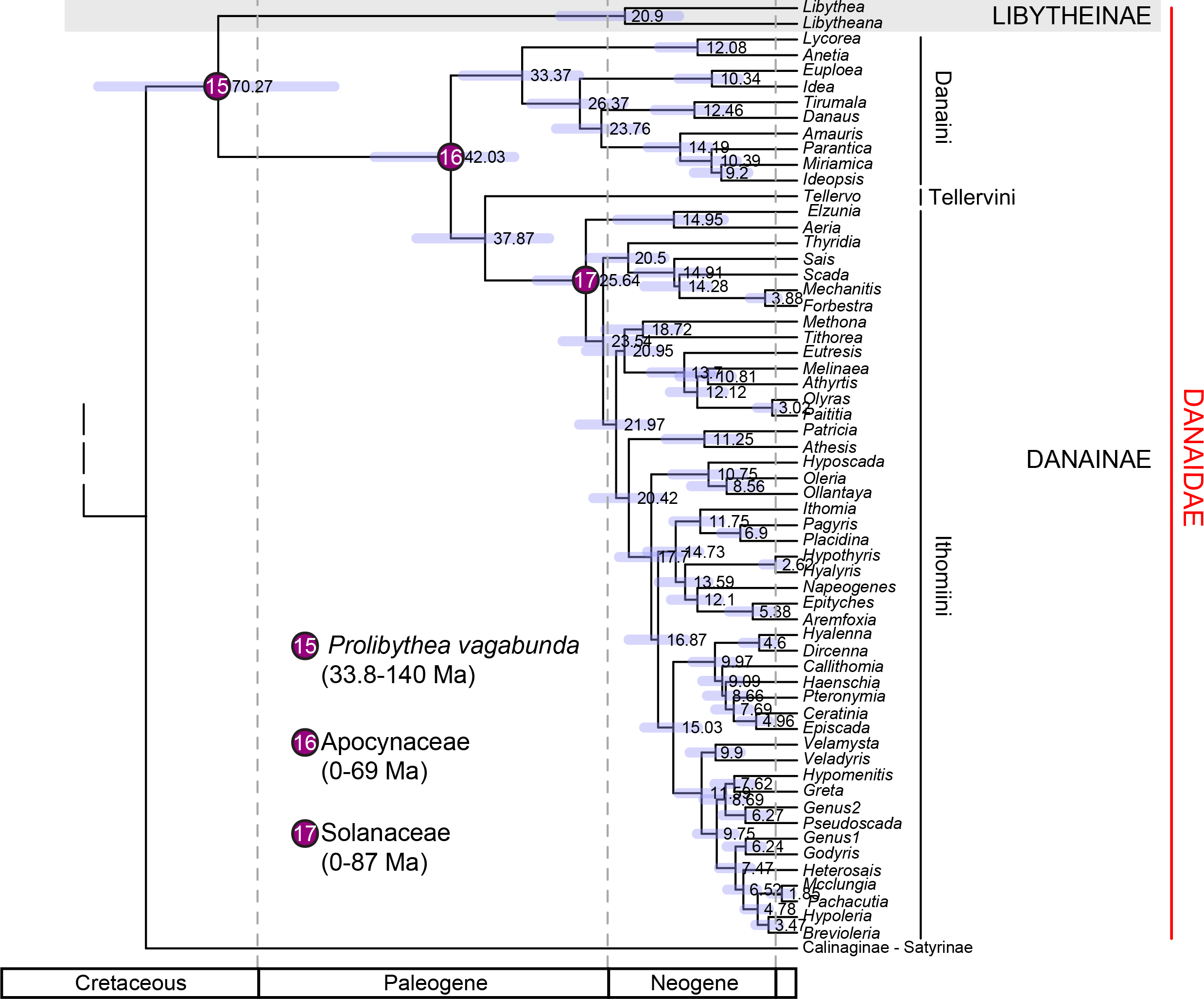
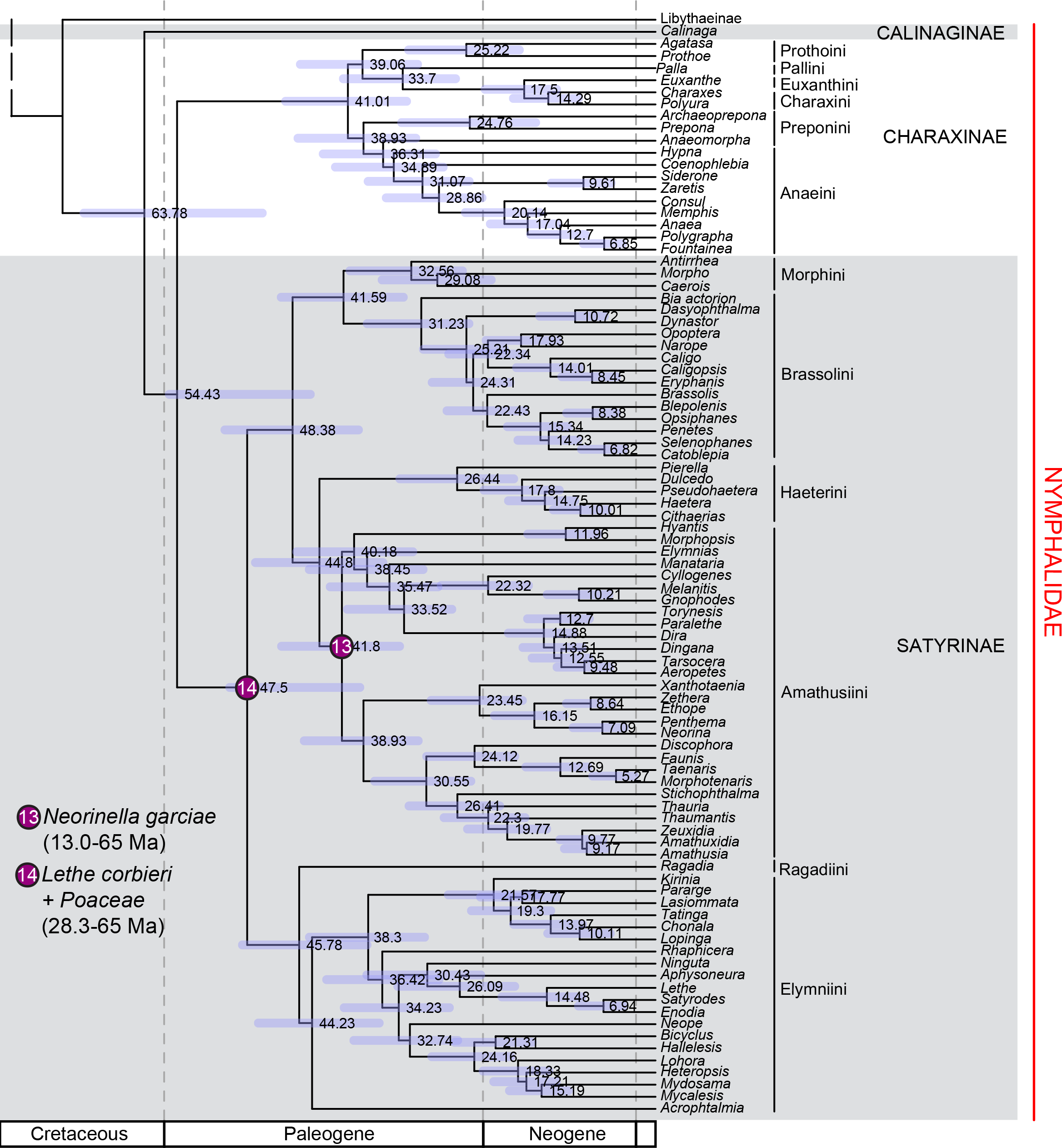
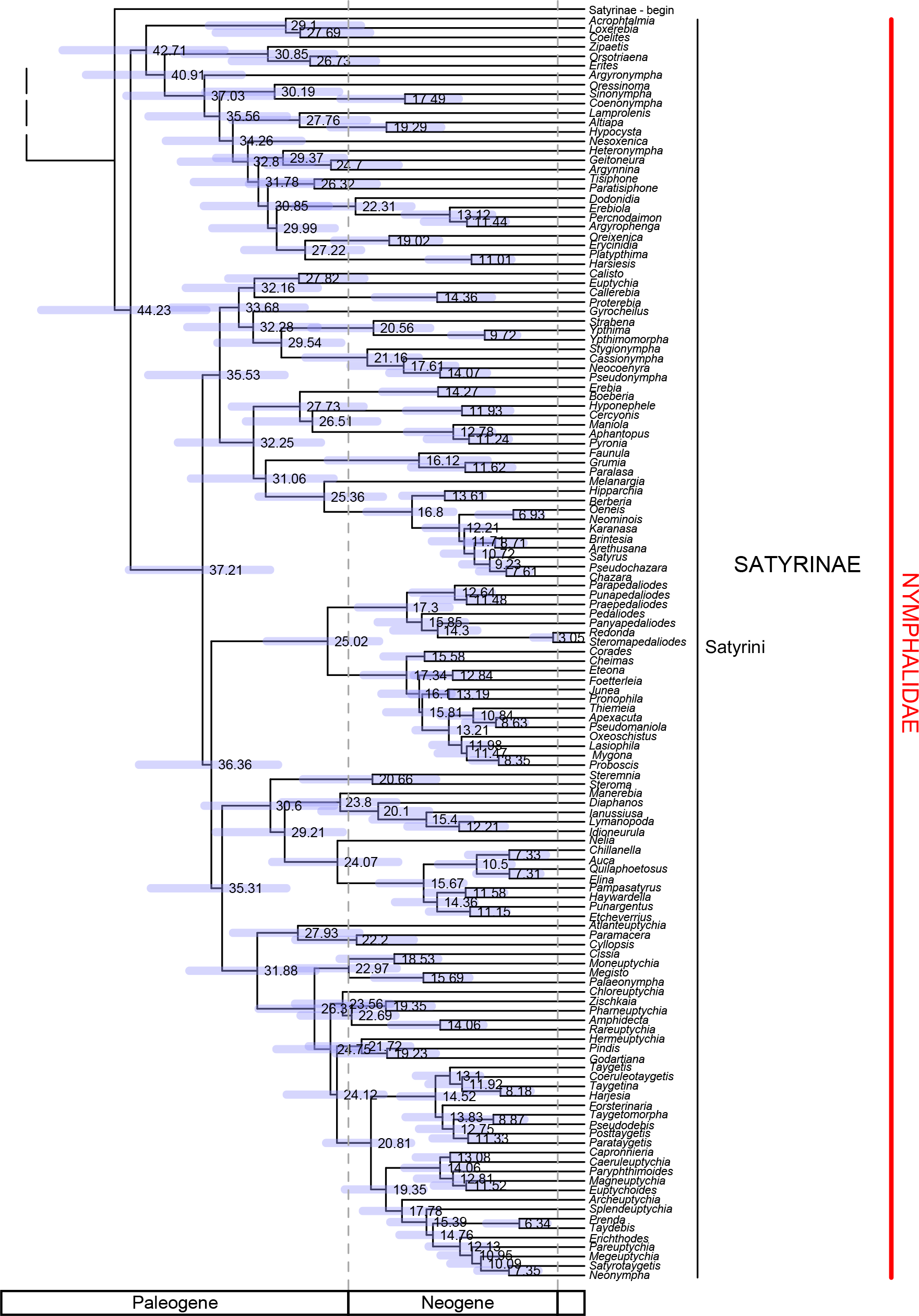
Time-calibrated tree obtained from the core analysis. a) The relationships and age estimates among the subfamilies of Papilionoidea. b) The relationships and age estimates among the genera across the different families. Age estimates are indicated at the nodes (Ma). Node bars represent the 95% credibility intervals.

**TABLE 1.** (a) Fossil calibration points used to calibrate the tree as a minimum age for the *Clade calibrated*. Unless stated otherwise, the fossil calibrations were placed at the stem of the clade calibrated. *Lower* and *upper* values indicate the prior truncation for both the uniform and exponential priors. The 140 Ma year upper truncation corresponds to the age of Angiosperms from Magallón et al. 2015. A different upper truncation value results from a fossil prior interacting with a host plant prior placed at the same node or a lower node. *Mean* and *offset* are parameter values for the exponential prior distribution. (b) Host-plant clades used to calibrate the tree as a maximum age for the *Calibrated node*. Host plant calibrations were placed at the crown of the clade calibrated. Ages from both Magallón et al. (2015) and Foster et al. (2017) are indicated.

| **Fossils** | **Clade calibrated** | **lower** | **upper** | **mean** | **offset** |
| --- | --- | --- | --- | --- | --- |
| *Doritites bosniaskii*<br>Rebel, 1898 | Papilionidae: Parnassiinae:<br>Luehdorfiini | 5.3 | 140 | 25 | 5.3 |
| *Dynamine alexae*<br>Peñalver & Grimaldi, 2006 | Nymphalidae: Biblidinae:<br>*Dynamine* | 15.9 | 89 | 20 | 15.9 |
| *Lethe corbieri*<br>Nel, Nel & Balme, 1993 | Nymphalidae: Satyrinae:<br>Satyrini | 28.3 | 65 | 25 | 28.3 |
| *Mylothrites pluto*<br>Heer, 1849 | Pieridae:<br>Coliadinae+Pierinae | 15.9 | 100 | 50 | 15.9 |
| *Neorinella garciae*<br>Martins-Neto et al., 1993 | Crown of Amathusiini | 23.0 | 65 | 20 | 23.0 |
| *Pamphilites abdita*<br>Scudder, 1875 | Hesperiidae: Hesperinae | 23.0 | 140 | 30 | 23.0 |
| *Prolibythea vagabunda*<br>Scudder, 1889 | Nymphalidae: Libytheinae | 33.8 | 140 | 40.0 | 33.8 |
| *Protocoeliades kristenseni*<br>de Jong, 2016 | Hesperiidae: Coeliadinae | 55.6 | 140 | 35 | 55.6 |
| *Thaites ruminiana*<br>Scudder, 1875 | Papilionidae: Parnassiinae:<br>Parnassiini | 23.0 | 140 | 25 | 23.0 |
| *Theope* sp | Riodinidae: Riodininae:<br>Nymphidiini: *Theope* | 15.9 | 140 | 25 | 15.9 |
| *Voltinia dramba*<br>Hall, Robinson & Harvey, 2004 | Riodinidae: Riodininae:<br>Mesosemiini: *Voltinea* | 15.9 | 140 | 30 | 15.9 |
| *Vanessa amerindica*<br>Miller & Brown, 1989 | Nymphalidae:<br>Nymphalinae: Nymphalini | 33.8 | 140 | 30 | 33.8 |
| *Doxocopa wilmattae*<br>Cockerell, 1907 | Nymphalidae:<br>Nymphalinae+Biblidinae+<br>Limenitidinae+Apaturinae |  | Not used |  |  |
| *Praepapilio colorado*<br>Durdin & Rose, 1978 | Papilionidae |  | Not used |  |  |

b)
| **Host plant clade** | **Clade calibrated** | **Magallón et al. 2015** | **Foster et al. 2017** |
| --- | --- | --- | --- |
| Angiospermae | root | 140 | 252 |
| Poaceae | Hesperiidae: Hesperinae | 65 | 112 |
| Poaceae | Nymphalidae: Satyrinae | 65 | 112 |
| Fabaceae | Pieridae | 100 | 123 |
| Brassicaceae | Pieridae: Pierinae | 103 | 97 |
| Rubiaceae | Riodinidae: Leucochimona+Mesophtalma+<br>Mesosemia+Perophtalma<br>+Semomesia | 87 | 85 |
| Apocynaceae | Nymphalidae: Danainae | 69 | 85 |
| Solanaceae | Nymphalidae: Ithomiini | 87 | 68 |
| Euphorbiaceae | Nymphalidae: Biblidinae | 89 | 104 |
| Sapindaceae | Nymphalidae: Biblidinae: Epiphilini+<br>Callicorini | 87 | 91 |

When a fossil was assigned to a clade, we calibrated the stem age of this clade, specifically the time of divergence from its sister clade, instead of the crown age or the first divergence event recorded in the phylogeny. As a consequence of this choice, we removed two of the 14 fossils. We did not use *Praepapilio colorado* Durden & Rose, 1978 (Papilionidae, 48.4 Ma) nor the less well-preserved *Praepapilio gracilis* Durden & Rose, 1978 (Papilionidae) of the same age because its position at the root of the tree was uninformative given the presence of the 55.6 million years old *Protocoeliades kristenseni* de Jong, 2016 placed at the crown of the Hesperiidae. For similar reasons, we did not use *Doxocopa wilmattae* Cockerell, 1907 (Nymphalinae+Biblidinae+Limenitidinae+Apaturinae, 33.8 Ma) because its position was uninformative given the presence of *Vanessa amerindica* Miller & Brown, 1989 of the same age but placed lower in the tree.

Host plant calibrations – Butterflies are well known for their strict relationships with specific groups of plants used by their larvae. Such associations have previously been suggested as evidence for coevolution (Ehrlich & Raven, 1964, Janz & Nylin 1998, Nylin & Janz 1999). In the present study, we selected nine calibration points based on known information of host plant specificity by butterflies since the large revision of Ackery (1988) (see also Beccalloni et al. 2008 for Neotropical species), and revised for those host plant records listed as having spurious or occasional records (AVLF unpublished data). Host plant clades used by single genera or a small group of recently-derived genera were discarded, such as the use of Aristolochiaceae by Troidini. In these cases the butterflies clearly are much more recent than their associated plant clades, and consequently do not contribute relevant time information to the tree. The ages of each plant group were defined as maximum ages for the respective nodes (Table 1). For all host plant maximum constraints we used the estimate from Magallón et al. (2015) using the upper boundary of the 95% credibility interval of the stem age of the host plant clade. We also constrained the root of the Papilionoidea with a maximum age corresponding to the crown age of angiosperms from Magallón et al. (2015). The host plant calibrations were placed at the crown of the butterfly clades as a conservative approach since we do not know when the host plant shift occurred on the stem branch. However, we assume that the diversification of the clade could not have begun earlier than the origin of the host plant family.

### Analyses Overview

Given computational limitations for such a large dataset, we adopted the following procedure (details given below). We ran PartitionFinder v. 1.1 (Lanfear et al. 2012) to identify the best partition scheme. Using this result we performed a maximum likelihood analysis to obtain a tree topology. This tree topology was transformed into a time-calibrated ultrametric tree and used thereafter as a fixed topology and starting tree in all our dating analyses. Branch lengths were estimated using BEAST v. 1.8.3 (Drummond et al. 2012) with a simpler partitioning scheme, a birth-death tree prior, lognormal relaxed molecular clocks, and a combination of minimum (fossils) and maximum (host plants) constraints for which all were set with uniform priors. This constituted the core analysis. We then performed additional analyses to test the robustness of our results to (i) different subsets of fossil constraints, (ii) the prior distribution of fossil constraints, (iii) a different estimate for host-plant ages, (iv) a Yule tree prior, (v) a reduced taxon sampling, and (vi) the addition of a mitochondrial gene fragment.

### Core Analysis

Tree topology – We started by running PartitionFinder v. 1.1 (Lanfear et al. 2012) on the concatenated dataset, allowing all possible combinations of codon positions of all genes. Substitution models were restricted to a GTR+G model and branch lengths were linked. We then performed a maximum likelihood analysis using RAxML v8 (Stamatakis 2006) using the best partitioning scheme identified by PartitionFinder and 1000 ultrafast bootstraps. The resulting tree was set as a fixed topology for the dating analyses. To do so, the tree was transformed into a time-calibrated ultrametric tree using the package *ape* (Paradis et al. 2004) and all minimum and maximum constraints in order to obtain a starting tree suitable for BEAST analyses.

Time tree – We used BEAST v. 1.8.3 (Drummond et al. 2012) to perform our time-calibration analysis. Given the size of our dataset, we reduced the number of partitions in our dating analysis to three partitions, each partition being one codon position of all genes pooled together. Substitution rate for each partition was modeled by GTR+G and an uncorrelated lognormal relaxed molecular clock. We used a Birth-Death process as branching process prior. In order to have a fixed topology we turned off the topology operators in BEAUTi and we specified the topology obtained with RAxML made ultrametric with the *ape* package.

Setting the priors for calibration points is always an important matter of discussion. Non-uniform priors are often used, yet in the majority of studies the choice of parameters defining the shape of the prior distribution is not justified (Warnock et al. 2012). For the core analysis we followed a conservative approach – considering that fossils only provide a minimum age, while host plant calibrations only provide a maximum age for the nodes they were assigned to – and we used uniform prior distributions for all calibration points (Table 1). When a node was calibrated with fossil information, the distribution ranged from the estimated age of the fossil to the age of angiosperm origin (extracted from Magallón et al. 2015). When a node was calibrated using host-plant age, the prior distribution ranged from 0 (present) to the age of the host plant clade origin. When a node was calibrated with both types of information, the distribution ranged for the age of the fossil to the age of host plant clade origin. We also used a uniform prior for the tree root height, ranging between the oldest fossil used in the analysis and the age of angiosperm origin. Host plant calibrations, as well as the origin of angiosperms were extracted from Magallón et al. (2015), using the upper boundary of the 95% credibility interval of the stem age of the host plant clade. Our choice of combining (1) uniform prior distributions, (2) fossil calibration of stem nodes, (3) the oldest stem age of the host plant clades and (4) host plant calibration of crown nodes has important implications. On the one hand these choices are the most conservative options, cautiously using the information given by each type of calibration point and taking into account uncertainty surrounding the information used. On the other hand, they are also the least informative.

We performed four independent runs of 30 million generations, sampling every 30 000 generations. We checked for a satisfactory convergence of the different runs using Tracer v. 1.6.0 (Rambaut et al. 2014) and the effective sample size values in combination. Using LogCombiner v. 1.8.3 (Drummond et al. 2012), we combined the posterior distributions of trees from the three runs, discarding the first 100 trees (10% burn-in) of each run. Using TreeAnnotator v. 1.8.3 (Drummond et al. 2012) we extracted the median and the 95% credibility interval of the posterior distribution of node ages.

### Alternative Analyses

We tested the effect of making alternative choices along the core analysis on our estimates of divergence times. Unless stated otherwise, we made only one modification at a time; all other parameters remained identical to that described for the core analysis. We performed at least two independent runs of 30 million generations per alternative parameter set and more if convergence was not reached.

Different subset of fossils – We aimed at testing whether using only a fraction of the fossil information affected the estimation of divergence times and whether the position of calibrations (close to the root or close to the tips) also changed the results. Thus, we divided our set of fossil constraints into two subsets depending on their position in the tree. One subset included fossil calibration points assigned at a deep level in tree (hereafter: higher-level fossils): *Lethe, Mylothrites, Neorinella, Pamphilites, Prolibythea, Protocoeliades* and *Vanessa* (Table 1). The other subset included fossil calibration points close to the tips of our phylogeny (hereafter: lower-level fossils): *Doritites, Thaites, Dynamine, Theope* and *Voltinia* (Table 1). In both cases the full set of maximum constraints was used. We performed one analysis for each subset.

Exponential fossil priors – In the core analysis we used uniform distributions for calibration points, which is a conservative option but also the least informative. As an alternative, we designed exponential priors for fossil calibration points. Exponential priors use the age of a fossil as minimum age for the node it has been assigned to, but also assume that the probability for the age of the node decreases exponentially as time increases. In BEAUTi, we set the offset of exponential distributions with the age of the fossil. The distribution was truncated at the maximum age used in the uniform priors. The shape of the exponential distribution is controlled by a mean parameter, which has to be arbitrarily chosen by the users. The choice of mean parameter can be found in Table 1. Priors for host plant calibration points were not changed (i.e., uniform priors).

Alternative host plant ages –The origin and timing of diversification of angiosperms is controversial. While the oldest undisputed fossil of Angiospermae is from the early Cretaceous (136 Ma, Brenner 1996), most divergence time estimations based on molecular clocks have inferred a much older origin. In the core analysis, we chose to use host plant ages derived from the tree of angiosperms time-calibrated by Magallón et al. (2015), who imposed a constraint on the origin of angiosperms based on this fossil information. They found a crown age for angiosperms of ~ 140 Ma. As an alternative consistent with an older origin of angiosperms we used ages recently inferred by Foster et al. (2017), who recovered a crown age of angiosperms of ~ 209 Ma. All maximum constraints were replaced by those inferred by Foster et al. (2017). The origin of angiosperms was used as a maximum constraint was set to the upper boundary of the 95% credibility interval of the crown age of the angiosperms i.e., 252.8 Ma. Because the posterior distributions of node ages for this analysis were very skewed, we extracted the median of the distribution, the 95 % credibility interval and the mode of the kernel density estimate of nodes using the R package *hdrcde*. For comparison, we also estimated the mode of posterior distributions for the core analysis and all alternative tests.

Yule branching process prior – Condamine et al. (2015) showed that the prior for the tree growth can a have a great impact on the estimated divergence times. In the core analysis we used a Birth–Death prior, which models the tree formation with a constant rate of lineage speciation and a constant rate of lineage extinction. As an alternative, we used a Yule prior, which involved a constant rate of speciation and no extinction to assess whether age estimates changed or not.

Reduced dataset – In our core analysis, we chose to maximize the taxon sampling – increasing the number of lineages – which increased the fraction of missing data in the molecular dataset. We tested whether increasing the molecular dataset completion to the detriment of taxon sampling changed the results. In this reduced dataset, we included all the genera for which a specific minimum number of genes were available. The missing data in the molecular dataset are not uniformly distributed across the tree; for example, Lycaenidae have more missing data than the Nymphalidae. Therefore, a different cut-off value was chosen for each family in order to keep a good representation of the major groups (Papilionidae: 5 genes, Hedylidae: 8 genes, Hesperiidae: 9 genes, Pieridae: 8 genes, Lycaenidae: 4 genes, Riodinidae: 8 genes, Nymphalidae: 9 genes). In order to allow assignment of all fossils to the same place as in the core analysis, nine taxa having a number of genes below the cut-off value had to be added. We ended up with a dataset reduced to only 364 taxa instead of 994 in the core analysis. Accordingly, the fraction of missing data decreased from 39.5% in the core analysis to 21.4%. Given this important modification of the dataset we generated a new topology with RAxML, which was then calibrated identically to the core analysis.

Mitochondrial gene fragment – We tested whether adding mitochondrial information in the dataset would affect our results. To do so, we added the cytochrome-oxydase-subunit 1 gene to the molecular dataset. Given the conflicting signal in Hesperiidae between nuclear and mitochondrial information (Sahoo et al. 2016), the COI was not added to the Hesperiidae. We performed a new RAxML analysis in order to obtain a new topology. This new tree was calibrated with BEAST identically to the core analysis, with one difference. The mitochondrial gene was added as two partitions separated from the nuclear partitions: the first and second positions of COI were pulled together and the third position had its own partition. Therefore this analysis had five partitions.

### Comparing Prior and Posterior Distributions

When performing a Bayesian analysis, comparing prior and posterior parameter distributions can be informative about the amount of information contained by our data compared to the influence of prior information. As exemplified by Brown & Smith (2017), such a comparison can shed light on the discrepancies observed in the fossil record and the divergence times estimated from a time-calibrated molecular clock. It may also help to disentangle the effect of interaction among calibration points. For each calibrated node we can compare the user-designed prior distribution (e.g., uniform distributions in the case of the core analysis), the marginal prior distribution that is the result of the interaction between the user priors and the tree prior, and the posterior distribution that is the distribution after observing the data. For the core analysis, the two different subsets of fossils and the alternative host plant ages analyses were re-run without any data to sample from the marginal prior. In each case we performed two independent runs of 50 million generations, sampling every 50 000 generations. The results were visualized with Tracer. When necessary, we performed an additional run. Using LogCombiner, the runs were combined after deleting the first 10% as burn-in. The results of the analyses with and without the molecular dataset were imported into R (R Development Core Team 2008) and for each calibrated node as well as the root height we compared the kernel density estimates of the marginal prior and the posterior distributions (R package *hdrcde*).

### Comparison with Previous Studies

For the root of all Papilionoidea and the seven families we compared the estimates obtained in the core analysis to previous studies that also used fossil information.

## RESULTS

### Core Analysis

The core analysis performed with BEAST used the full set of fossils and host plant constraints from Magallón et al. (2015) on the topology found with RAxML. This analysis resulted in a root estimate for all Papilionoidea of 107.6 Ma (Fig. 1, Supplementary Material S2). The 95% credibility interval of the posterior distribution ranged from 88.5 to 129.5 Ma. The lineage leading to Papilionidae diverged first at the root of Papilionoidea and the crown age of Papilionidae was inferred to be 68.4 Ma (95%CI=53.5–84.3). Hedylidae and Hesperiidae diverged from Pieridae–Lycaenidae–Riodinidae–Nymphalidae at 106.5 Ma (95%CI=88.0–127.2) and diverged from each other at 99.2 Ma (95%CI=80.7–119.2). The crown age of the sampled Hedylidae was 32.8 Ma (95%CI=23.4–43.6) and crown age of Hesperiidae was 65.2 Ma (95%CI=55.8–78.1). Pieridae diverged from Lycaenidae–Riodinidae–Nymphalidae at 101.1 Ma (95%CI=83.0–120.3) and extant lineages started diversifying around 76.9 Ma (95%CI=63.1–92.4). Lycaenidae and Riodinidae diverged from Nymphalidae at 97.4 Ma (95%CI=80.4–116.5) and diverged from each other at 87.8 Ma (95%CI=73.2–106.1). The crown age of Lycaenidae was 71.0 Ma (95%CI=57.2–85.2) and crown age of Riodinidae was 73.4 Ma (95%CI=60.3–88.1). Finally, the crown age of Nymphalidae was inferred to be 82.0 Ma (95%CI=68.1– 98.3).

### Alternative Analyses

In most cases the seven alternative parameters tested yielded very similar results (Fig. 2, Supplementary Material S3-S8). Reducing the number of taxa in order to decrease the fraction of missing data, using higher-level calibration points only, or using a Yule process tree prior (instead of a Birth–Death prior), gave virtually identical results as the core analysis above. Using only lower-level fossil constraints (close to the tips of the phylogeny) resulted in the youngest estimates of all alternative runs, with a crown age of Papilionoidea of 94.5 Ma (mode=83.8, 95%CI=67.8–126.6). Using exponential fossil priors mainly resulted in a narrower credibility interval, while the mode and median age estimates were only 7–8 million years younger than the core analysis mode estimate (Fig. 2, Supplementary Material S6). Adding mitochondrial information also lead to a 7–8 million-year younger estimate for the crown age of Papilionoidea, but the credibility interval remained comparable to the core analysis (Supplementary Material S7). Finally, using a hypothesis of older host plant ages extracted from Foster et al. (2017), we obtained the greatest difference. The upper boundary of the credibility interval largely shifted toward much older ages (95%C I=88.5–167.2) as well as the median (119.5 Ma). The posterior distribution was, however, very skewed, with a mode of 101.0 Ma, and converged to the same age as the core analysis (Fig. 2, Supplementary Material S8).

**Figure 2.**
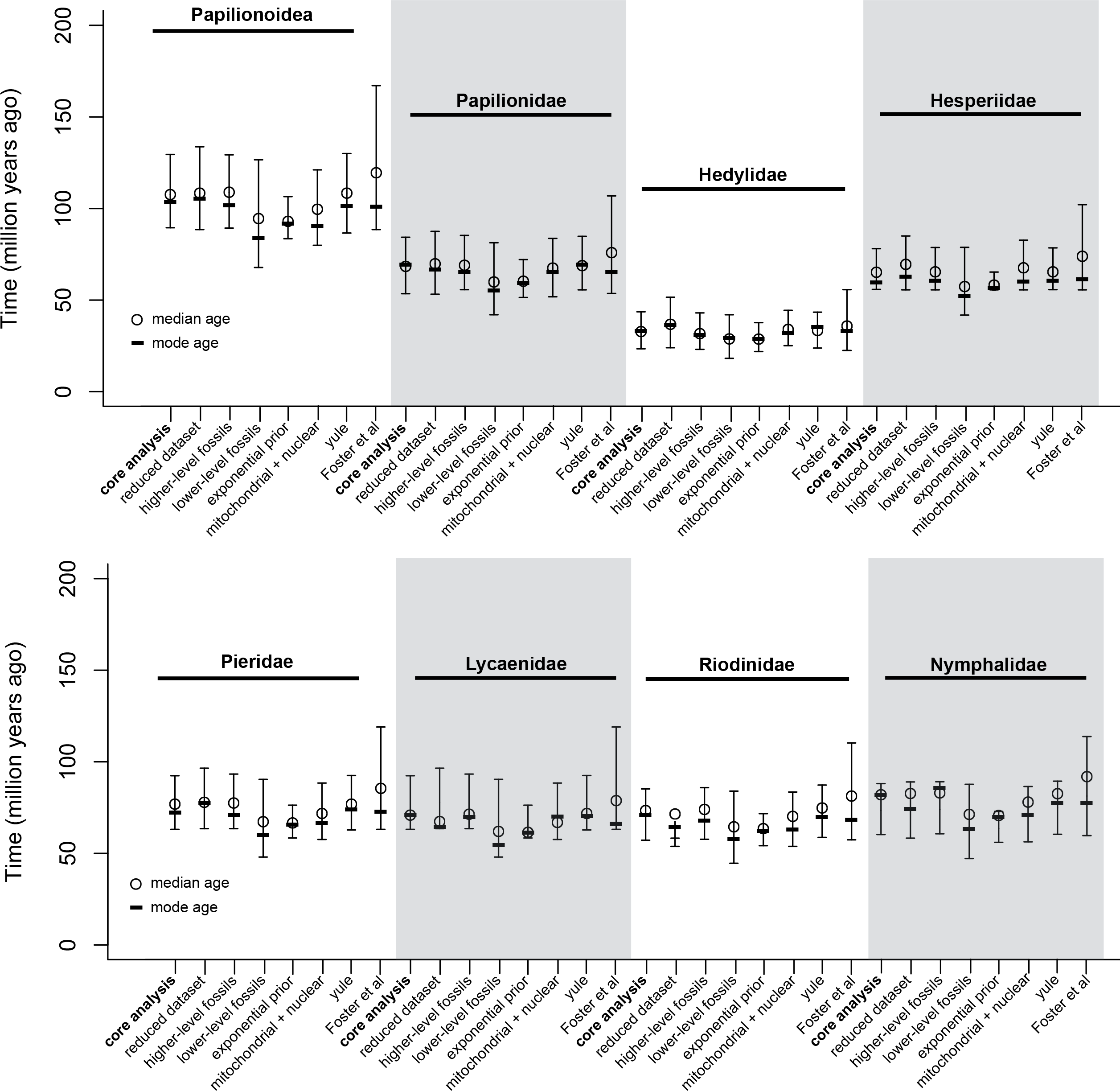
>Comparison of node age estimates for the root of Papilionoidea and the seven families between the core analysis and the seven alternative analyses. Mode, median and 95% credibility interval are presented.

These variations for the root age among different alternative analyses were recovered for the ages of the different subfamilies. For example, all lower-level fossils always led to younger estimates while older ages from Foster et al. (2017) always led to older estimates (Fig. 2).

### Comparing Prior and Posterior Distributions

We compared the posterior distributions to the marginal prior distributions for the different calibrated nodes in the core analysis. We set all fossil and host plant constraints with uniform prior distributions as we considered this as the most conservative approach. However, it is important to note that the marginal prior distributions at these nodes, which result from the interactions between all calibration priors and tree prior, are not uniform (Fig. 3).

**Figure 3.**
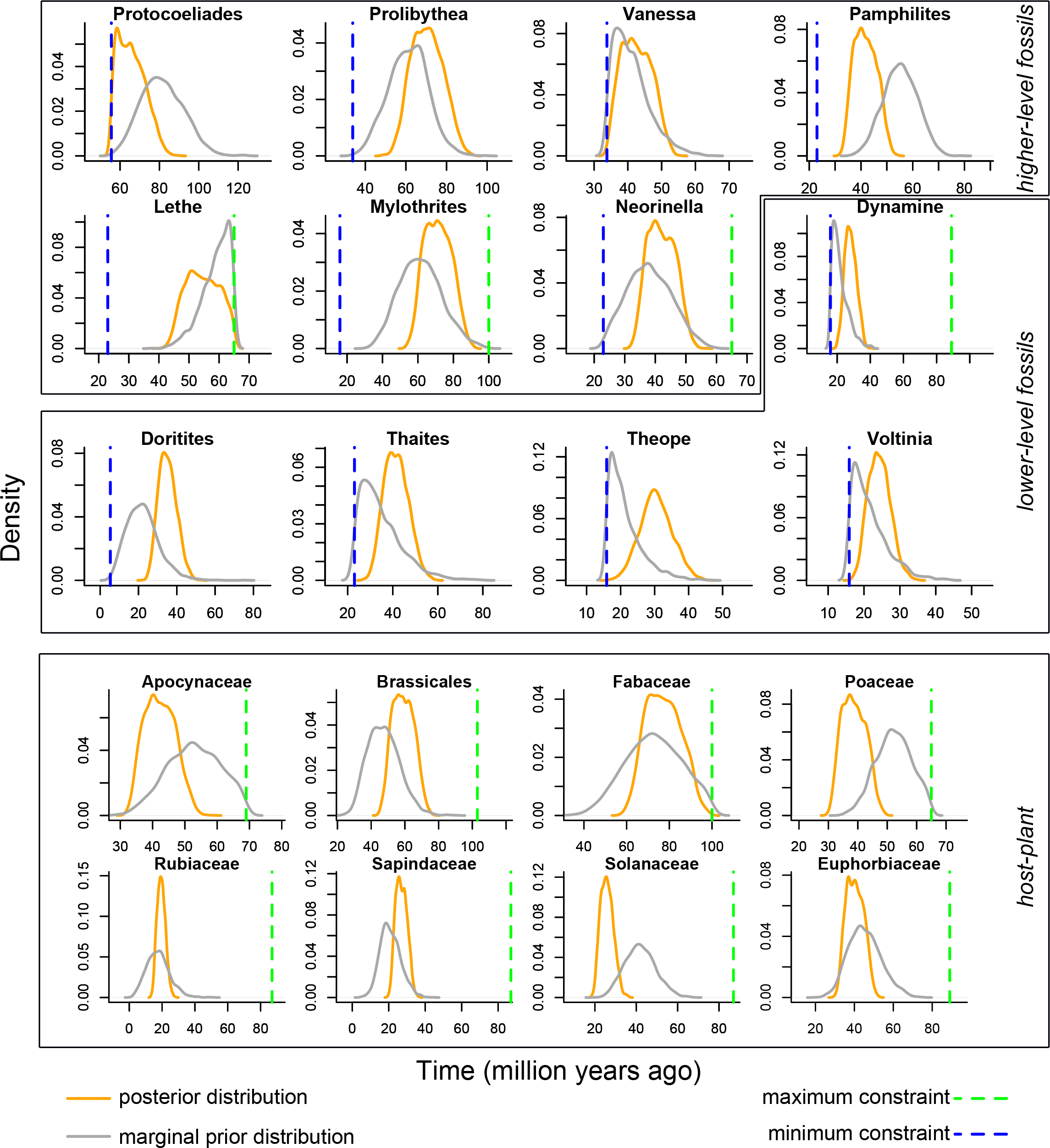
Marginal prior (grey) and posterior distributions (orange) for the nodes calibrated in the core analysis. Blue dashed lines represent minimum boundaries; green dashed lines represent maximum boundaries.

Across all calibrated node points, many of them showed shifts of posterior distributions from the marginal priors, indicating that the results of the core analysis was not a simple outcome of our set of priors (Fig. 3). Interestingly, the nodes calibrated by *Doritites*, *Dynamine*, *Thaites*, *Theope* and *Voltinia*, which are all the fossils placed close to the tips of our phylogeny, tended to shift away from the minimum boundary, toward older ages than the marginal prior distribution.

Alternative analyses performed with only these lower-level fossils yielded the youngest tree for butterflies. This suggests that higher-level fossils bring important additional information, leading posterior distributions of lower-level nodes to shift away from the prior distributions in the core analysis.

The nodes calibrated with the higher-level fossils *Mylothrites, Prolibythea, Neorinella* and *Vanessa* showed posterior distributions largely overlapping with their marginal prior distributions. Many host plant calibrated points showed a shift from the marginal prior distribution (Fig. 3). In all cases, except the node also calibrated with the fossil *Lethe* the crown age of the butterfly clade inferred was much younger than the age of the corresponding host plant clade.

For the root of Papilionoidea, the marginal prior and posterior distributions largely overlapped in the core analysis, therefore not indicating whether our molecular dataset contained significant information about the root age or not. We also compared the posterior and the marginal prior distributions for alternative analyses performed with different subsets of fossil calibrations (Fig. 4). When using only higher-level fossils, the posterior distribution was almost identical to the core analysis, but the marginal prior slightly shifted from the marginal prior of the core analysis toward a younger age. The use of only lower-level fossils had more profound effects. In such a case, prior distributions of the core analysis and the lower-level fossils alternative completely overlapped. The posterior distribution, however, shifted toward younger ages, yielding the most recent estimate for the root age among all analyses (mean=94.5, mode=83.8, 95%CI=67.8–126.5). We also looked at the effect of using relaxed maximum ages (based on Foster et al. 2017). In this case, marginal prior distribution for the root age shifted to a mean of ~148 Ma (Fig. 4) and a credibility interval spanning 100 Ma (95%CI=99.9–205.8). The posterior distribution was very skewed, retaining a wider credibility interval than the core analysis (95%CI=88.5– 167.5), but significantly shifted from the prior distribution toward the posterior distribution of the core analysis (median=119.5, mode=101.0).

**Figure 4.**
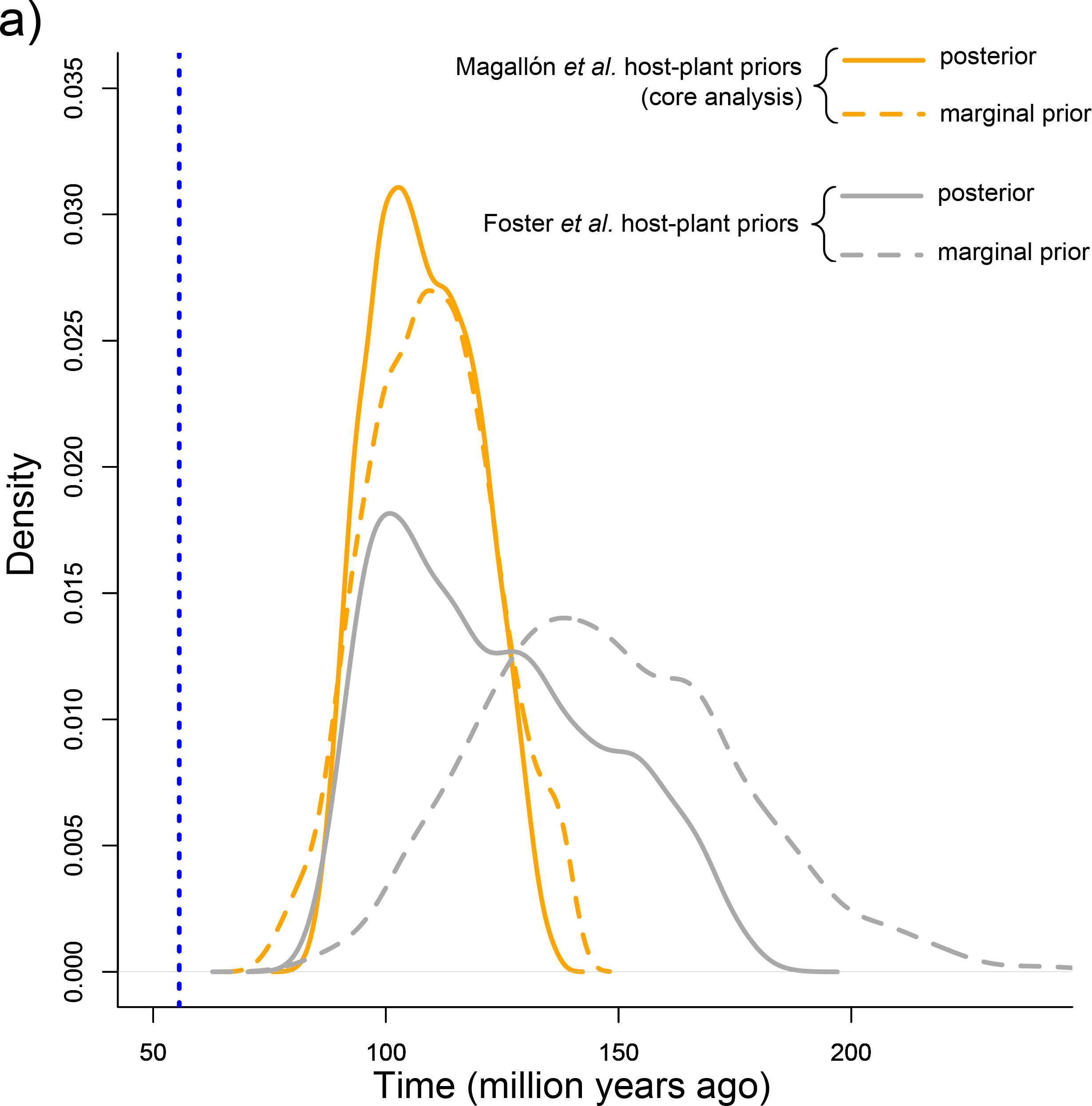
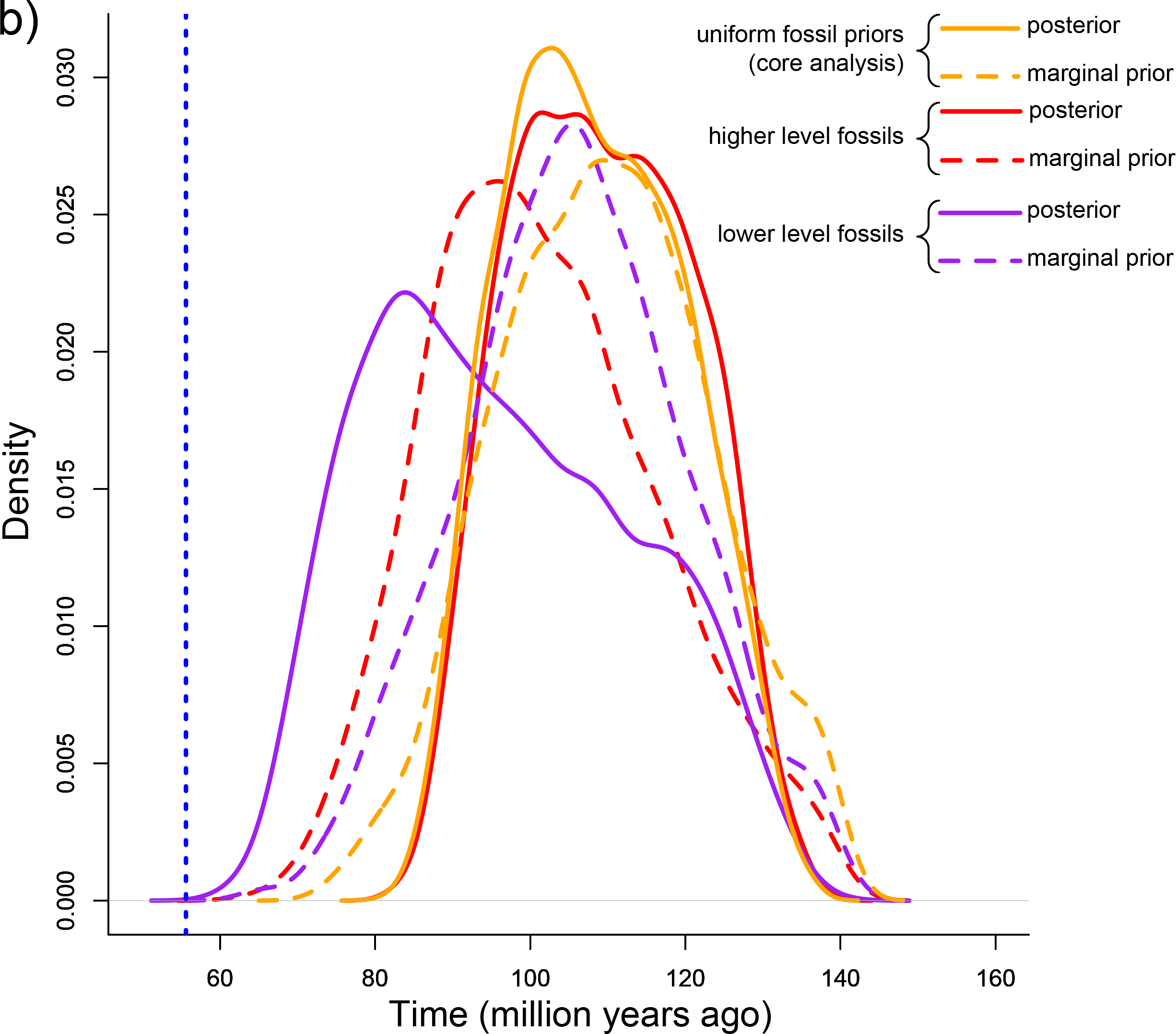
Marginal prior and posterior distributions for the root age in the core analysis using either a) alternative host-plant ages or b) alternative subsets of fossil calibrations.

### Comparison with Previous Studies

For the root of Papilionoidea, our estimate in the core analysis using the mode age of the distribution was very similar to Wahlberg et al. (2013) and Heikkilä et al. (2012), with a mean age estimate of 104.6 and 110.8 Ma, respectively (107.6 Ma in the core analysis, Fig. 5). For the crown age of families our estimates were often consistent with most of previous studies. For Papilionidae, our crown age estimate (68.4, 95%CI=53.5–84.3) was very similar to Wahlberg et al. (2013) and Heikkilä et al. (2012), while Condamine et al. (2012) in a study focusing primarily on this family found younger ages of about 15 million years. For Hedylidae, only the study by Heikkilä et al. (2012) had an estimate for the crown age, whose mean age was 45.3 Ma, which is older than our result (32.8, 95%CI=23.4–43.6). The age of Hesperiidae (65.2, 95%CI=55.8–78.1) was similar to Wahlberg et al. (2013) and Heikkilä et al. (2012), but much younger than Sahoo et al. (2017) with an estimate of 82 Ma. Pieridae is the family that showed highest variation in age estimates among different studies. Our estimate (76.9 Ma, 95%CI=63.1–92.4 Ma) falls in between the youngest estimate from Wahlberg et al. (2013), in which the credibility interval goes down to 39 Ma, and the oldest estimate from Braby et al. (2006), in which the oldest boundary of the credibility interval was 111.6 Ma. For Lycaenidae, which contain no fossils calibrations, the results between our core analysis (73.4, 95%CI=60.3–88.1), Wahlberg et al. (2013) and Heikkilä et al. (2012) were virtually identical. For the crown age of Riodinidae, our core analysis (70.9, 95%CI=57.2–85.2) gave identical results to Heikkilä et al. (2012). Espeland et al. (2015), in a study focusing specifically on this family found about 10 million–year older ages. Wahlberg et al. (2013), however, found a much younger estimate, about 20 Ma younger. For Nymphalidae, we have the greatest number of time calibrations, but they all tend to find very similar results. Our estimation (82.0, 95%CI=68.1–98.3) was very close to Wahlberg et al. (2013) and Heikkilä et al. (2012). This estimation was about 12 million years younger than the study by Wahlberg et al. (2009) focusing on Nymphalidae.

**Figure 5.**
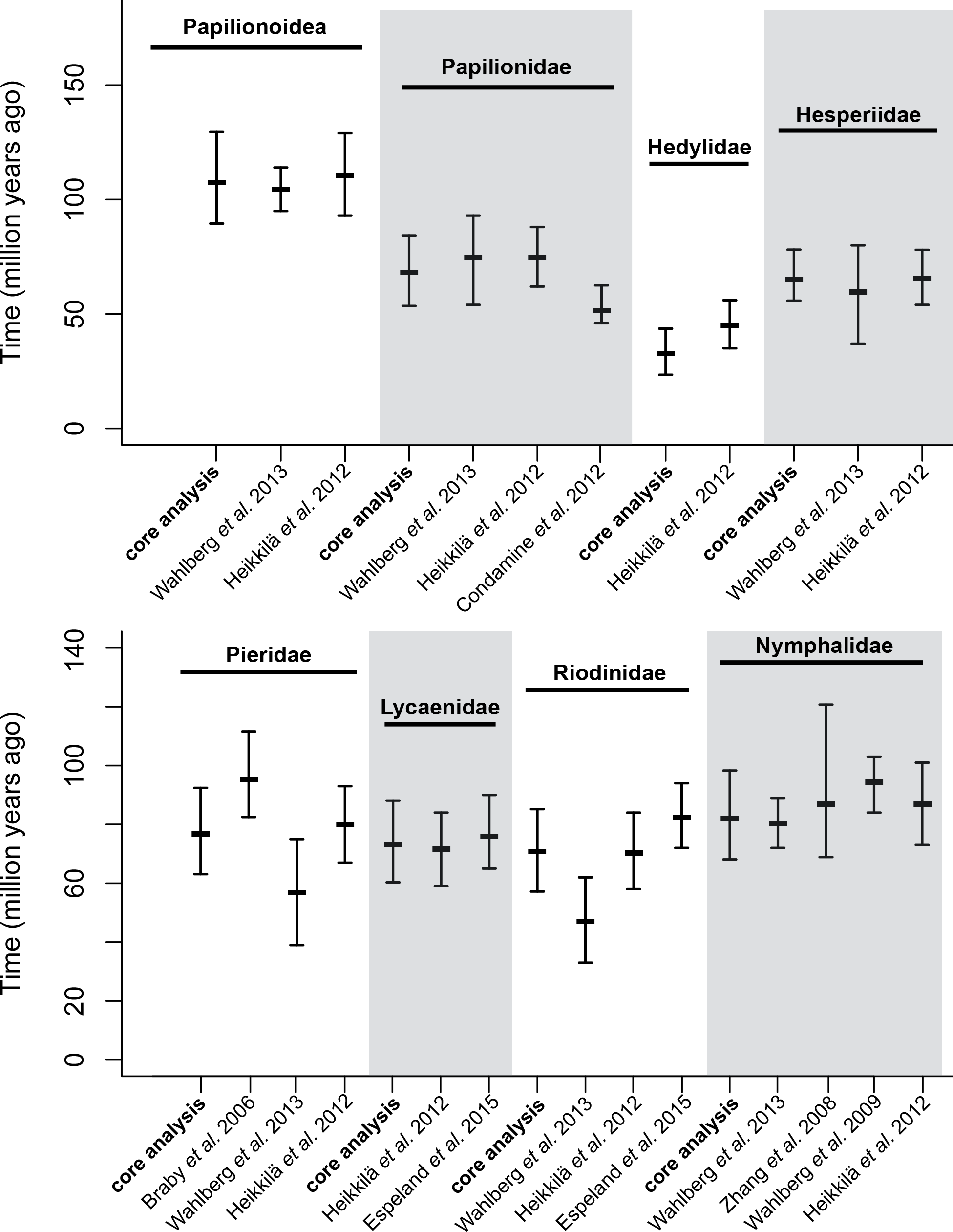
Comparison of node age estimates for the root of Papilionoidea and the seven families between this study (core analysis) and estimates from previous studies. Mode and 95%CI for the core analysis are presented. For the other studies the values reported in the original study are used.

## DISCUSSION

We generated a genus-level phylogeny of the superfamily Papilionoidea, including 994 taxa. Taking advantage of a recent revision of the lepidopteran fossil record we established a new set of 12 fossil calibration points, which were combined to 10 secondary calibrations from host plant ages.

### Fossils and Minimum Ages

In the core analysis we adopted a very conservative approach. This choice involves taking into account the uncertainty surrounding the information available for each calibration point, although at the expense of the amount of useful information available. For fossil constraints, this had two consequences. First, we calibrated the stem of the focal clade a fossil was assigned to, by calibrating the divergence from its sister group instead of the first divergence recorded in the phylogeny within the focal clade itself. Calibrating the crown age of the focal clade – meaning that we assume that the fossil is “nested” within the clade – may lead to an overestimation of the crown age. Such would be the case if lineages are undersampled at the root, or if extinction occurred, or if the fossil belongs to a lineage that actually diverges somewhere along the stem. Calibrating a higher node with the age of the fossil, which involves loss of some information, is considered to be the best way to avoid these problems. Second, we used uniform prior distributions bounded by the age of the fossil and the age of angiosperms. We considered that fossils provide only a minimum age for a node, a condition that is especially exacerbated by the exceptionally poor fossil record of Lepidoptera in general (Labandeira and Sepkoski, 1993) and Papilionoidea in particular (Sohn et al. 2015) when compared to the four other major hyperdiverse insect lineages (Coleoptera, Hymenoptera, Diptera and Hemiptera). Prior expectation on the age of the node cannot be modeled more accurately without additional information. However, the marginal priors resulting from the interactions among the different priors actually strongly differ from this assumption.

### Higher-versus Lower-Level Calibrations

Generally, favoring multiple calibrations placed at various positions in a tree instead of a single or few calibrations, seem to produce more reliable estimates of molecular clocks (Conroy & Van Tuinen 2003, Smith & Peterson 2002, Soltis et al. 2002, Duchêne et al. 2014). Calibrations distributed across a tree may allow a better estimation of substitution rates and their pattern of variation among lineages (Duchêne et al. 2014), and improve age estimates in cases of taxon undersampling (Linder et al. 2005).

Calibrations placed at deep levels in the tree are usually favored (Sauquet 2012, Hug & Roger 2007) over calibrations at lower levels for better capturing the overall genetic variation (Duchêne et al. 2014). Yet, deep calibrations also tend to underestimate the mean substitution rate and lead to an overestimation of shallow nodes, referred to as “tree extension” by Phillips (2009). For the butterflies, we investigated the consequences of using different subsets of fossil calibrations according to their positions in the tree (higher *versus* lower-level calibrations), compared to the full set of fossil constraints. With a subset of fossils placed only at higher levels in the phylogeny we obtained results similar to the full set of fossils in the core analysis, either at the deep nodes or shallow nodes, indicating no tree extension effect. This effect may also indicate that the lower level calibration points that are close to the tips are uninformative, and when included in the core analysis, do not affect the timescale but clearly affected the priors (see below).

Alternatively, lower-level calibrations can lead to an overestimation of the mean substitution rate across the tree, thereby underestimating the timescale (Phillips 2009). Interestingly, when only a subset of fossils were used and placed close to the tips, it led to the youngest estimates, including the credibility intervals. This potentially indicates an effect of mean substitution rate overestimation. Also, we noticed in the core analysis that the nodes calibrated by *Protocoeliades* and *Vanessa* (two deep node constraints) showed posterior distributions abutting against the minimum boundaries defined by the age of the fossils, therefore preventing the tree (or at least these nodes) to be younger.

### Host Plants and Maximum Ages

For calibration points constrained by the age of the host plant group, we considered that only the crown of the focal clade could be assigned confidently to the host plant group, as the stem or part of the stem could be older than the host plant (the host plant shift would be happening somewhere along the stem). Support arises from molecular biological and paleobiological evidence that the establishment of specialized insect-herbivore associations can considerably postdate the origins of their hosts, as in a Bayesian analysis of 100 species of leaf-mining *Phyllonorycter* moths (Lepidoptera: Gracillariidae) and their dicot angiosperm hosts (Lopez-Vaamonde et al. 2006). Relying on host plant ages for calibrating a butterfly tree is questionable while the timing of the divergence of angiosperms is still highly controversial (e.g. Magallón et al. 2015, Foster et al. 2017). As a result, first we calibrated our tree using the oldest boundary of 95% CI of the stem age of a host plant clade. This allowed us to take into account the uncertainty surrounding the timing of the first appearance of the host plant but consequently, it also relaxed the prior hypothesis for the calibrations. Secondly, we compared two alternative timescales for the angiosperms: a paleontological estimate, which infers an Early Cretaceous origin of angiosperms (Magallón et al. 2015), and a molecular clock estimate that we extracted from Foster et al. (2017), which infers a stem age for angiosperms during the Early Triassic about 100 million years older. These two alternative scenarios affected the size of the credibility intervals and the shape of the posterior distributions. For the crown of Papilionoidea, the upper boundary of the 95%CI was ~37 million years older when using the molecular clock estimate. However, the shape of the distribution was very asymmetrical, with a mode of the distribution very close to the core analysis (101.0 Ma), suggesting that the estimation for the age of the root still concentrated around the same ages. Using the hypothesis of an Early Triassic origin of angiosperms implied very permissive priors toward old ages, which are most likely responsible for the very wide credibility intervals and asymmetrical posterior distributions recovered in the alternative analysis of using ages from Foster et al. (2017). Therefore, it is tempting to use the time-scale inferred using Magallón et al. (2015)’s ages of angiosperms, as it greatly narrows down the uncertainty surrounding butterfly ages, and aligns more realistically with the fossil angiosperm record. However, as long as there is no consensus on the timing of angiosperm diversification there is no reason to favor one or the other.

### Priors and Posterior Distributions

We compared the marginal priors to the posterior distributions for different analyses for the root of Papilionoidea and for the different calibration points in the core analysis. We found several calibration points showing a substantial shift of posterior distribution. This indicates that our age estimates are not entirely driven by the set of constraints, but instead that the molecular dataset brings additional information about the age of the calibrated nodes. An interesting pattern we found in the core analysis is the consistent trend of posterior distributions of the lower-level calibrated nodes to shift toward older ages than the priors. Meanwhile, some higher-level node calibrations shifted toward younger ages than the prior but most of them largely overlapped with their prior distribution. Consequently, posterior estimates tend to contract the middle part of tree compared to the prior estimates.

There are at least three reasons for the anomalous gap between the earliest fossil papilionoid occurring at 55.6 Ma and its corresponding Bayesian median age of 110 Ma, that represents a doubling of the lineage duration. First, it long has been known that the lepidopteran fossil record is extremely poor when compared to the far more densely and abundantly occurring fossils of the four other hyperdiverse, major insect lineages of Hemiptera, Coleoptera, Diptera and Hymenoptera (Labandeira and Sepkoski, 1993). Second, particularly large-bodied apoditrysians such as Papilionoidea, have even a poorer fossil record than other Lepidoptera in general, particularly as they bear a fragile body habitus not amenable to preservation. Additionally, as external feeders papilionoids lack a distinctive, identifiable trace fossil record such as leaf mines, galls and cases (Sohn et al. 2015). Third, there are very few productive terrestrial compression or amber deposits spanning the Upper Cretaceous, from 100 Ma to the Cretaceous-Paleogene boundary of 66.0 Ma, and the part of the Paleogene Period from 66.0 Ma to the earliest papilionoid fossil of 55.6 Ma (Labandeira, 2014; Sohn et al., 2015). Some of these deposits have recorded very rare small moth fossils, but to date no papilionoid, or for that matter, other large lepidopteran taxa such as saturniids or pyraloids have been found.

The root of the tree was only calibrated with the oldest fossil in our dataset, a 55.6 million-year-old papilionoid, and the crown age of the angiosperms. However, the prior distribution for the root in the core analysis clearly excluded an origin of butterflies close to 55.6 Ma, but rather a distribution centered on a median of 110 and ranging between 86.4 and 136.2 Ma. The posterior distribution for the root in the core analysis largely overlapped with the prior. However, when we used alternative ages for the angiosperms (older ages), the marginal prior for the root shifted to substantially older ages. Nevertheless, the posterior distribution showed a significant shift toward younger ages, albeit highly skewed, toward ages similar to the core analysis. This suggests that our estimate of the root age in the core analysis is not simply driven by our set of priors, even if we do not actually observe a shift between marginal prior and posterior distributions.

We observed some differences in prior and posterior distributions at the root when considering only subsets of fossils. When using only the subset of higher-level fossils, the marginal prior for the root showed very little difference from the core analysis prior and the posterior distributions completely overlapped. When using the subset of lower-level fossils the marginal prior remained similar to the core analysis but the posterior distribution showed a substantial shift toward younger ages, yielding the youngest estimation of the age of Papilionoidea among all our analyses. As such, it seems that the choice of fossils did not change the prior estimation of the root, but the posterior distribution was largely influenced by higher-level fossils. As we suggested earlier, lower-level fossils only may be overestimating the mean substitution rate across the tree, and therefore underestimating the time scale, while the implementation of higher-level fossils seems to be correcting for this.

### Timescale of Butterflies Revisited

We propose a new estimate for the timing of diversification of butterflies, based on an unprecedented set of fossil and host-plant calibrations. We estimated the origin of butterflies between 89.5 and 129.5 Ma, the median of this posterior distribution being 107.6 Ma, which corresponds to the Early Cretaceous–Late Cretaceous boundary interval. The result of our core analysis for the root is very close to previous estimates by Wahlberg et al. (2013) and Heikkilä et al. (2012). The comparisons of alternative analyses, the prior and posterior distributions showed that this result is robust to almost all the choices made throughout the core analysis and that our molecular dataset contains significant information in addition to the time constraints. This estimation means that there is a 45 million-year-long gap between the oldest known butterfly fossil and the molecular clock estimate. Accordingly, as Brown & Smith (2017) stated for the case of angiosperms, we do not know whether a larger molecular dataset – implying potentially more information for estimating the molecular clock– would allow the root to become younger. Alternatively, the fossil record for butterflies is so sparse that an intervening fossil gap is very likely. Besides, the fossil *Protocoeliades kristenseni*, which is 55.6 Ma can be assigned confidently to the crown of the family Hesperiidae and the stem of Coeliadinae well within the Papilionoidea. For angiosperms, a very rich fossil record is available compared to butterflies (e.g., Magallón et al. (2015), which used 137 fossils to calibrate a phylogeny of angiosperms), rendering the absence of angiosperms, either as pollen or macrofossils, that are older than 136 Ma much more puzzling.

## ACKNOWLEDGMENTS

This is contribution 366 of the Evolution of Terrestrial Ecosystems consortium at the National Museum of Natural History, in Washington D.C. NW acknowledges funding form the Swedish Research Council (Grant No. 2015-04441) and from the Department of Biology, Lund University. AVLF thanks CNPq (grant 303834/2015-3), National Science Foundation (DEB-1256742) and FAPESP (grant 2011/50225-3). This publication is part of the RedeLep (Rede Nacional de Pesquisa e Conservação de Lepidópteros) SISBIOTABrasil/CNPq (563332/2010-7). MH gratefully acknowledges funding from a Peter Buck Postdoctoral Stipend, Smithsonian Institution National Museum of Natural History.

## SUPPORTING INFORMATION

S1. List of taxa and Genbank accession codes.

S2. Tree obtained from the core analysis. Node ages are the median of node age posterior distributions.

S3. Tree obtained from the reduced dataset. Node ages are the median of node age posterior distributions.

S4. Tree obtained when using only higher-level fossil calibrations. Node ages are the median of node age posterior distributions.

S5. Tree obtained when using only lower-level fossil calibrations. Node ages are the median of node age posterior distributions.

S6. Tree obtained when using exponential fossil calibration priors. Node ages are the median of node age posterior distributions.

S7. Tree obtained when adding a mitochondrial gene fragment. Node ages are the median of node age posterior distributions.

S8. Tree obtained when using the host-plant ages obtained from Foster et al. (2017).

In S8a node ages are the median of node age posterior distributions, while in S8b the node ages are the mode the mode of the kernel density estimate of the posterior distribution.

